# Dectin-3 recognizes cryptococcal glucuronoxylomannan to initiate host defense against cryptococcosis

**DOI:** 10.1101/281790

**Authors:** Hua-Rong Huang, Fan Li, Hua Han, Quan-Zhen Lv, Xia Xu, Ning Li, Shunchun Wang, Jin-Fu Xu, Xin-Ming Jia

## Abstract

*Cryptococcus neoformans* and *Cryptococcus gattii* cause life-threatening meningoencephalitis and pneumonia in immunosuppressed and immunocompetent individuals. Given the structural differences of major polysaccharide glucuronoxylomannan (GXM) between *C. neoformans* and *C. gattii*, it remains unclear that how innate immune system recognizes GXM. Here, we report that C-type lectin receptor Dectin-3 (MCL encoded by Clec4d) is a direct receptor for GXMs from *C. neoformans* serotype AD (*C.n*-AD) and *C. gattii* serotype B (*C.g*-B). GXMs from *C.n*-AD and *C.g*-B activated both NF-κB and ERK pathways to induce the pro-inflammatory cytokine production, whereas it was completely abolished due to deficiency of Dectin-3 or its downstream adaptor protein CARD9. Upon pulmonary *C.n*-AD and *C.g*-B infection, Dectin-3- and CARD9-deficient mice were highly susceptible and showed augmented lung injury due to impairment of alveolar macrophage accumulation and killing activities. These results demonstrate that Dectin-3 contributes to host immunity against *Cryptococcus* infection through selectively recognizingGXM.

## The Paper Explained

### Problem

The incidence of cryptococcosis caused by *C. neoformans* and *C. gattii* is nowadays particularly dramatic in developing countries. *C. neoformans* and *C. gattii* were subdivided serologically into five serotypes based on their major capsular polysaccharide glucuronoxylomannan (GXM). In detail, *C. neoformans* consists of serotypes A, D, and AD hybrid, and *C. gattii* consists of serotypes B and C. Recent studies show that innate immune response mediated by TLR including TLR2 and TLR4, or C-type lectin receptor including Dectin-1, Dectin-2 and Dectin-3 is dispensable for pulmonary infection with *C. neoformans* serotype A. Therefore, it remains poorly known that how host innate immune system recognizes *C. neoformans* and *C. gattii* to elicit antifungal immunity against their infections.

### Results

In the present study, we have extensively characterized the role of Dectin-3 and its downstream adaptor protein CARD9 against pulmonary cryptococcosis. We found that, when infection with *C. neoformans* serotype AD and *C. gattii* serotype B strains, Dectin-3- and CARD9-deficient mice were highly susceptible and showed augmented lung injury. More importantly, we showed that Dectin-3 directly recognized GXMs extracted from *C. neoformans* serotype AD and *C. gattii* serotype B strains to activate CARD9-mediated NF-κB and ERK pathways for initiating anti-fungal host responses against pulmonary cryptococcosis.

### Impact

Our study provides the first biological and genetic evidence demonstrating that Dectin-3 recognizes GXM of *C. neoformans* serotype AD and *C. gattii* serotype B to initiate host defense against cryptococcosis.

## Introduction

The saprophytic, encapsulated fungal pathogens *Cryptococcus neoformans* and *C. gattii* can cause life-threatening meningoencephalitis and pneumonia in immunocompromised and immunocompetent individuals, which is called cryptococcosis[1]. *C. neoformans* has been classified into three serotypes including A, D, and AD hybrid whereas serotypes B and C have been recognized as a separate species called *C. gattii* based on antigenic differences in the polysaccharide capsules of the fungus[2-4]. *C. neoformans* serotype A (*C.n*-A) is the most clinically prevalent in immunocompromised individuals including HIV patients, renal transplant recipients and those undergoing immunosuppressive therapy[5], whereas *C. neoformans* serotype D (*C.n*-D) is mostly found in Europe but has a sporadic global distribution[6]. A recent survey in Europe revealed that 19% of human infections are caused by *C. neoformans* serotype AD (*C.n*-AD), which is a hybrid of serotype A and D strains[7]. In contrast, infection by *C. gattii* including serotype B and C (*C.g*-B and *C.g*-C) is much less common in immunocompromised patients but is thought to be more virulent than *C. neoformans* and causes disseminated infections even in healthy hosts[8]. Previously, it was thought that *C. gattii* infections were restricted to tropical and subtropical regions[9], but the emergence of the outbreak events due to *C. gattii* infections in temperate areas of North America suggest a more global distribution of this yeast[10,11].

Both *C*. *neoformans* and *C. gattii* are found ubiquitously in soil and other niches, and inhalation of yeast or desiccated basidiospores into the lungs is extremely common, causing about 70% of children with pulmonary infections in urban environments[12]. Host immune cells including alveolar macrophages and dendritic cells recognize pathogen associated molecular patterns (PAMPs) *via* pattern recognition receptors (PRRs), which elicit the host defense response. C-type lectin receptors (CLRs), a PRR-recognizing PAMP composed of polysaccharides, have garnered the attention of many investigators in the study of host defense against fungal infection[13,14]. The genetic deficiency of Dectin-1, are presentative CLR recognizing β1,3-glucans of *Candida albicans* yeast and *Aspergillus fumigatus* conidia[15,16], did not influence the clearance of *C.n*-A pulmonary infections [17]. Dectin-2 is known to recognize α-1,2-mannans of *C. albicans* and *A. fumigatus* hyphae, lipophilic and hydrophilic components of Malassezia, mannose-capped lipoarabinomannan of *Mycobacterium tuberculosis* and unknown components of non-capsular *C.n*-A to trigger the production of various cytokines and chemokines, including pro-inflammatory Th1, Th17, and also Th2 cytokines[18-22]. Our earlier studies show that Dectin-3 (also called CLECSF8, MCL, or Clec4d) can recognize α-1,2-mannans of *C. albicans* hyphae and trehalose 6,6’-dimycolate (TDM) of *M. tuberculosis* [23,24]. However, a recent study shows that Dectin-3 is dispensable for mediating protective immune responses against pulmonary *C.n*-A infection[25]. The genetic defect of the caspase recruitment domain family member 9 (CARD9), an adaptor protein that operates downstream of CLRs for activating NF-κB and extracellular signal-regulated protein kinase (ERK) pathways[26,27], confers susceptibilities to pulmonary *C.n*-A infection probably due to the reduced accumulation of IFN-Γ-expressing NK and memory Tcells[28].

Cryptococcal capsule is composed primarily of glucuronoxylomannan(GXM), which comprises more than 90% of the capsule’s polysaccharide mass[29]. The typical GXM consists of a linear (1→3)-α-D-mannopyranan bearing β-D-xylopyranosyl (Xylp), β-D-glucopyranosyluronic acid (GlcpA), and 6-O-acetyl substituents[30-32]. The disposition of the O-acetyl substituents is the major determinant of the antigenic activity observed among GXMs obtained from all serotypes (A, B, C, D, and AD)[33]. The ability of GXM to activate the Toll-like receptor (TLR)-mediated innate immune response has been reported in several studies [34-36]. In detail, GXM from *C.n*-A activates TLR4-mediated intracellular signaling [34], but its contribution to the global innate response against *C. neoformans* infections is limited [37,38]. GXM from *C.n*-A can also interact with TLR2 [34], which is believed to influence the response to cryptococcal infection [35]. A recent study shows that GXM from five cryptococcal serotypes were differentially recognized by TLR2/TLR1 and TLR2/TLR6 heterodimers[36]. Overall, most studies on the immunological functions of GXM have focused on the polysaccharide fractions from *C.n*-A isolates. Given the structural differences of GXM among the five serotypes, it also remains unknown that how host innate immune system differentially recognizes these GXMs.

In the present study, we show that Dectin-3 is a direct receptor for capsular GXM from *C.g*-B and *C.n*-AD, but not *C.n*-A, *C.n*-D and *C.g*-C. Furthermore, we demonstrate that Dectin-3 is essential for GXM-induced inflammatory responses through activating NF-κB and ERK pathways. Therefore, Dectin-3-deficient mice are highly sensitive to pulmonary *C.g*-B and *C.n*-AD infections due to impairment of alveolar macrophage activation.

## Results

### Dectin-3 recognizes the capsule of *C.g*-B to induce NF-κB and ERK-mediated pro-inflammation responses

In nature and under laboratory conditions, the capsule of *Cryptococcus* strains is relatively thin but can be mature to become thick during mammalian infection[39]. In this study, *C.g*-B strain ATCC32609 was prepared as thin- and thick-capsulated suspensions as described[39] and capsule induction was confirmed by India ink staining (**Figure 1A**). To explore whether Dectin-3 is required for *C.g*-B induced pro-inflammation responses, we stimulated bone marrow derived macrophages (BMDMs) from wild-type (WT, *Clec4d^+/+^*) and Dectin-3 deficient (*Clec4d^−/−^*) mice with thin- and thick-capsulated *C.g*-B, and found that both thin- and thick-capsulated *C.g*-B strongly induced nuclear translocation of NF-κB (p65 subunit) (**Figure 1B**), together with IκBα phosphorylation and degradation (**Figure 1C**). In contrast, Dectin-3 deficiency completely impaired the *C.g*-B-induced NF-κBnuclear translocation (**Figure 1B**) and IκBα phosphorylation and degradation (**Figure 1C**). Moreover, both thin- and thick-capsulated *C.g*-B triggered sustained phosphorylation of ERK in WT BMDMs (**Figure 1C**), but Dectin-3 deficiency dramatically impaired *C.g*-B-induced ERK activation (**Figure 1C**). Together, these results indicate that Dectin-3 is critical for *C.g*-B-induced activation of NF-κB and ERK pathways. Furthermore, we found that secretion of pro-inflammation cytokines including TNF-α and IL-6 were increased in WT BMDMs when stimulated with thin- and thick-capsulated *C.g*-B, whereas Dectin-3 deficiency significantly impaired these responses (**Figure 1D**). However, *C.g*-B-induced levels of IL-12p40 and IL-1β were comparable in WT and Dectin-3 deficient BMDMs (**Figure 1D**). These results suggest that Dectin-3 is essential for *C.g*-B-induced NF-κB and ERK-mediated pro-inflammationresponses.

**Figure 1.**
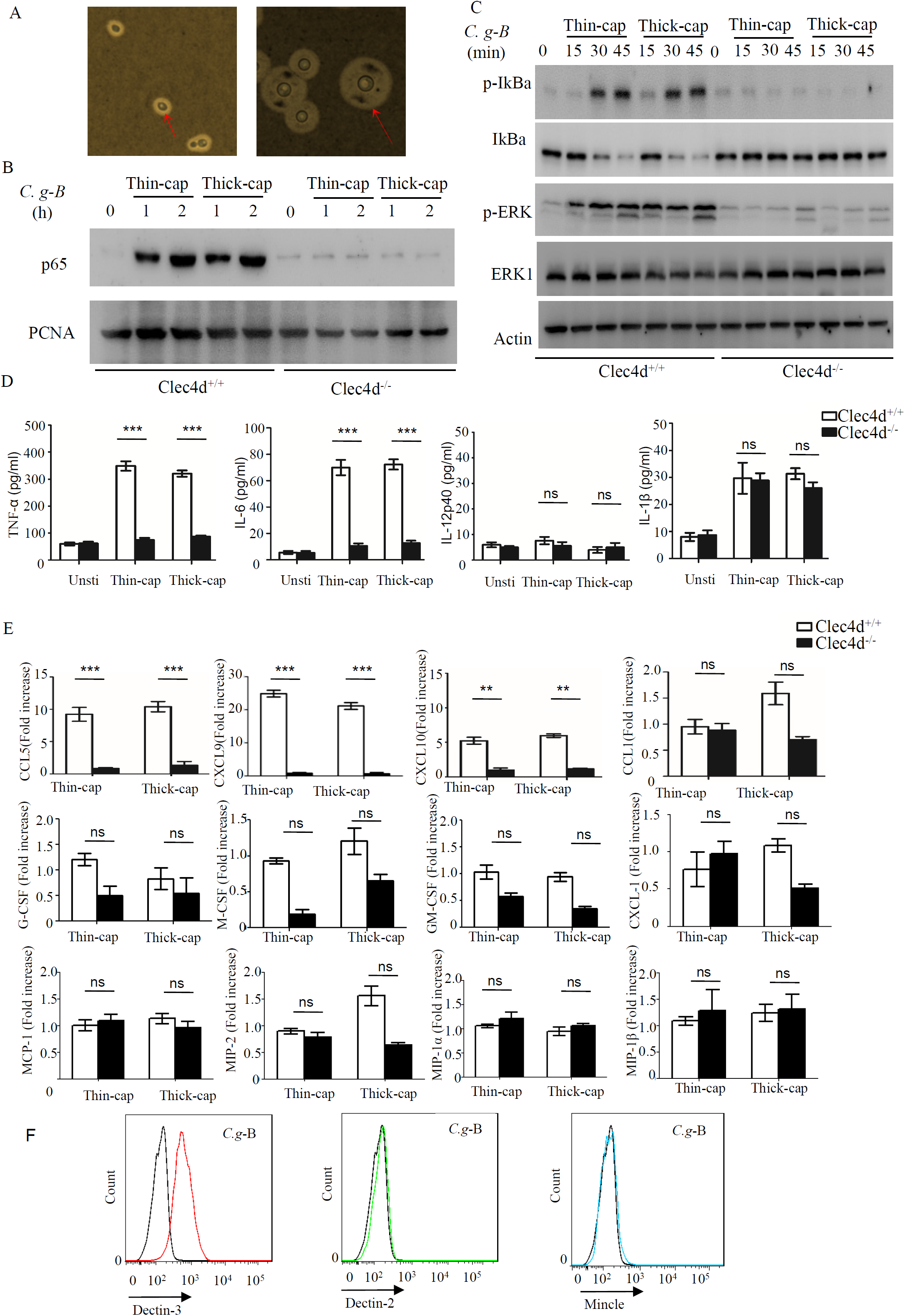
Dectin-3 is essential for *C. gattii*-B-induced NF-κB and ERK-mediated pro-inflammation responses. **(A)** *C.g-* B strain ATCC32609was cultured with thin-capsule form in YPD medium at 30°C and induced into thick-capsule form in RPMI-1640 medium plus 10% FBS at 37°C with 5% CO_2_ for 16h. Arrows indicates capsule of *C.g-* B strain. **(B and C)** protein phosphorylation **(C)** levels in BMDMs from WT and Dectin-3(Clec4d)-deficient mice, which were stimulated with thin- and thick-capsulated *C.g*-B strain (MOI=5) for indicated time. **(D)** ELISA results for indicated cytokines in WT and Dectin-3-deficient BMDMs, which were stimulated with thin- and thick-capsulated *C.g*-B strain (MOI=5) for 16h. Data are means ± SD of triplicate wells and are representative of three independent experiments;^***^p<0.001. **(E)** mRNA expression levels of indicated chemokines in WT and Dectin-3-deficient BMDMs, which were stimulated with thin- and thick-capsulated *C.g*-B strain (MOI=5) for 3h. **(F)** FACS assay of the binding of human Dectin-2-Fc, Dectin-3-Fc and Mincle-Fc fusion protein to *C.g-* B compared with a control human IgG Fc fusion protein (Control-Fc).

During cryptococcal infection, various chemokines are produced by alveolar monocytes to attract inflammatory cells and specific leukocyte subsets from blood to the site of infection. Here, we found that Dectin-3 deficiency in BMDMs strikingly impaired *C.g*-B-induced expression of CCL5/RANTES and of C-X-Cmotif chemokine ligand 9 (CXCL9) and CXCL10 (**Figure 1E**), which can attract activated T cells (particularly Th1 cells), dendritic cells, and monocytes by binding to specific CC chemokine receptors (CCR) on these cells, including CCR5 and CXCR3 [28]. However, there were slight influences of Dectin-3 deficiency on *C.g*-B-induced expression of CCL1, CXCL-1, macrophage inflammatory protein-1α (MIP-1α), MIP-1β, MIP-2, monocyte chemotactic protein-1 (MCP-1), granulocyte-macrophage colony-stimulating factor (GM-CSF), G-CSF and M-CSF in BMDMs (**Figure 1E**). These results suggest that Dectin-3 is also essential for *C.g*-B-induced NF-κB and ERK-mediated chemokine productions, resulting in the formation of granulomas and activation of adaptive immunity.

To explore whether Dectin-3 can bind the surface capsule of *C.g*-B, we performed a fluorescence-activated cell-sorting (FACS) screening assay for *C.g*-B binding using a panel of soluble receptor fusion constructs composed of the extracellular portion of Dectin-3, Dectin-2 and Mincle fused with the Fc fragment of human IgG1 antibody (**Figure S1**). As shown in **Figure 1F**, Dectin-3 was the only one of the soluble receptors that can bind the capsule of *C.g-* B and no binding was observed to Dectin-2 and Mincle. Together, these results indicate that Dectin-3 recognizes the capsule of *C.g*-B to initiate NF-κB and ERK-mediated pro-inflammation and chemo-attractive responses.

### Dectin-3 recognizes the capsule of *C.n*-AD to induce pro-inflammation responses

A recent study shows that Dectin-3 is not required for mediating protective immune responses against pulmonary *C.n*-A infection[25]. Here, we found that stimulation with *C.n*-A failed to induce the production of pro-inflammation cytokines including TNF-α, IL-6, IL-12p40 and IL-1β in WT and Dectin-3 deficient BMDMs (**Figure 2A**). However, stimulation with *C.n*-AD, *C.g*-C and *C.n*-D could induce large amounts of pro-inflammation cytokines including TNF-α and IL-12p40 in WT BMDMs whereas *C.n*-AD stimulation induced IL-6 and IL-1β production (**Figure 2A**). Of note, Dectin-3 deficiency significantly impaired the production of TNF-α, IL-6 and IL-12p40 in BMDMs when stimulated with both thin- and thick-capsulated *C.n*-AD, but not *C.g*-C and *C.n*-D (**Figure 2A**). As controls, Dectin-3 deficiency only affected TNF-α and IL-6 secretion in BMDMs when stimulated with another *C.g*-B strain WM179 (**Figure 2A**). Unexpectedly, Dectin-3 deficiency had no influence on the IL-1β production when stimulated with *C.n*-AD and *C.g*-B (**Figure 2A**). These results suggest that Dectin-3 is also critical for mediating pro-inflammation responses induced by *C.n*-AD, but not *C.n-* A, *C.n-* D and *C.g*-C.

**Figure 2.**
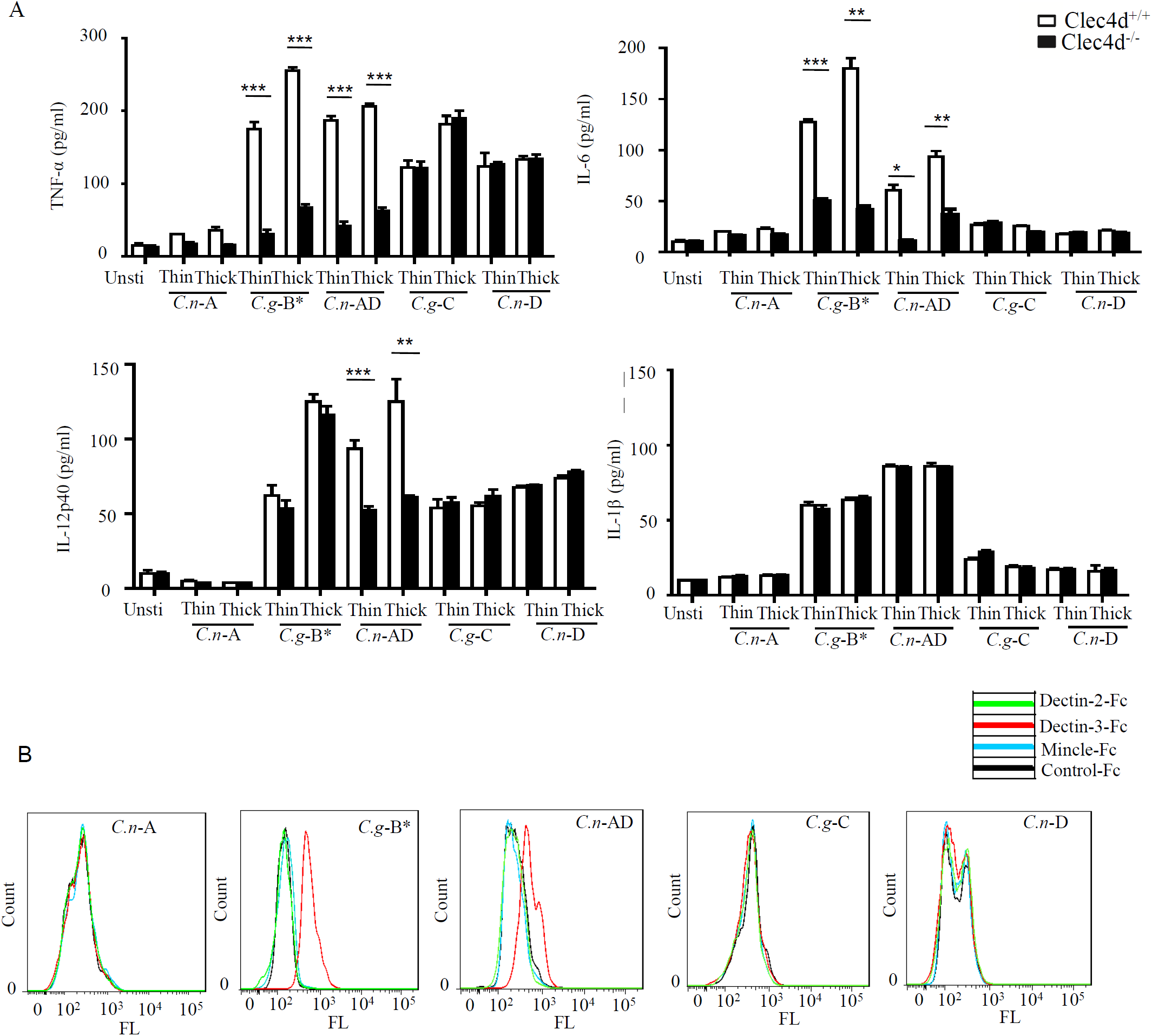
Dectin-3 is critical for NF-κB and ERK-mediated pro-inflammation responses induced by *C. neoformans*-AD. **(A)** ELISA results for indicated cytokines in WT and Dectin-3-deficient BMDMs, which were stimulated with thin- and thick-capsulated *C.n*-A strain H99, *C.g-* B strain WM179, *C.n-* AD strain WM628, *C.g-* C strain NIH312 and *C.n-* D strain WM629 (MOI=5) for 16h. Data are means±SD of triplicate wells and are representative of three independent experiments; **p<0.01,***p<0.001. **(B)** FACS assay of the binding of human Dectin-2-Fc, Dectin-3-Fc and Mincle-Fc fusion protein to *C.n-* A strain H99, *C.g-* B strain WM179, *C.n-* AD strain WM628, *C.g-* C strain NIH312 and *C.n-* D strain WM629 compared with a control human IgG Fc fusion protein (Control-Fc).

Furthermore, we performed FACS screening assay to examine the binding of Fc-fusion Dectin-2, Dectin-3 and Mincle to the surface capsule of *Cryptococcus* strains including *C.n-* A, *C.n-* D, *C.n*-AD and *C.g*-C, with another *C.g*-B strain WM179 as a control. As shown in **Figure 2B**, only Dectin-3 could bind the capsule of *C.n*-AD and *C.g-* B and no binding was observed to Dectin-2 and Mincle with the surface capsule of *Cryptococcus* strains. Together,these results indicate that Dectin-3 recognizes the capsule of *C.n*-AD and *C.g-* B to initiate the pro-inflammation responses.

### Dectin-3 directly recognizes GXM from *C.g*-B and *C.n*-AD

It has been well-documented that cryptococcal capsule is primarily composed of GXM[29]. We extracted extracellular polysaccharides (EPS) from *C.n*-A, *C.g*-B, *C.g*-C, *C.n*-D and *C.n*-AD as previously reported and the extract procedure was shown in **Figure S2A**. The extracted EPS fraction is a mixture of several polysaccharides, so we performed anion column chromatography to obtain major components for exploring their structural differences. Since the GXMs from *Cryptococcus* strains contained uronic acid[32,40], we performed reduction reactions to change glucuronic acid (GluA) into glucose (Glu) of EPS for GC-MS analysis (**Figure S2B** and **Figure 3A**). Monosaccharide composition analysis showed that the EPS from *C.n*-A, *C.g*-B, *C.g*-C, *C.n*-D and *C.n*-AD mainly contained Glu, xylose (Xyl), and mannose (Man) (**Figure S2B** and **Figure 3A**), indicating that EPS are primarily composed of GXM, a major cryptococcal capsule component. It has been reported that GXMs from *Cryptococcus* strains are all comprised of a core repeating unit to which (1→2)-linked and (1→4)-linked β-D-Xyl units are added in increments of one to four residues (**Figure 3B**)[32,40]. GXMs from *C.n*-A and *C.n*-D are mainly substituted with Xyl at C-2, whereas GXMs from *C.g*-B, *C.g*-C and *C.n*-AD are substituted at C-2 and at C-4 (**Figure 3B**). Analytical data showed that molar ratios of Glu/Xyl/Man in *C.n*-A, *C.g*-B, *C.g*-C, *C.n*-D and *C.n*-AD were calculated as 0.67:2:3, 0.6:3:3, 0.5:4:3, 0.55:1:3 and 0.58:2:3, respectively (**Figure S2C** and **Figure 3C**), indicating that the quantity and position of xylose residue is major structural difference of GXMs among *C.n*-A, *C.g*-B, *C.g*-C, *C.n*-D and *C.n*-AD, and these structural difference may determine the differential recognition of GXM from *C.n*-A, *C.g*-B, *C.g*-C, *C.n*-D and *C.n*-AD by host PRRs.

**Figure 3.**
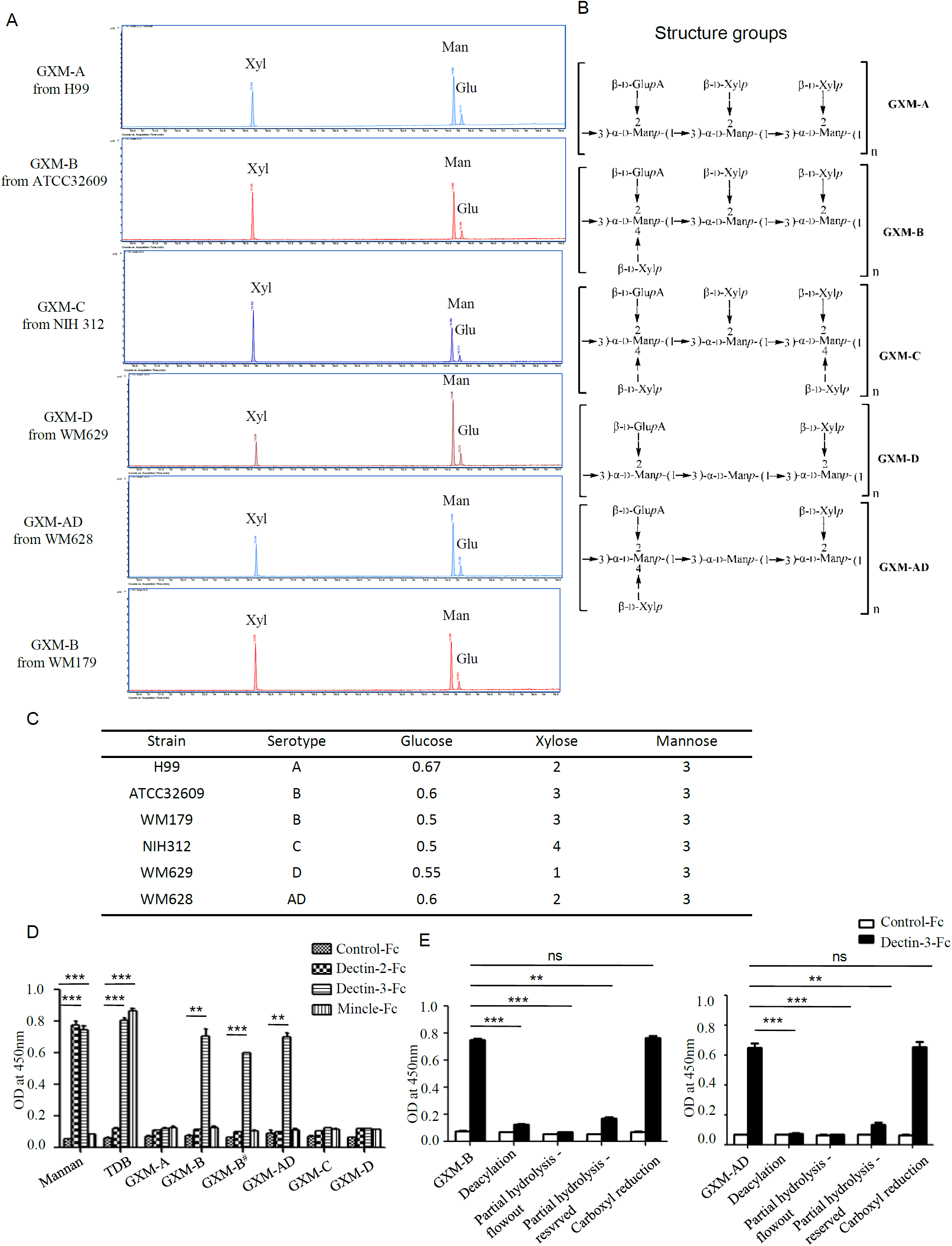
Dectin-3 directly recognizes GXM from *C.gattii*-B and *C.neoformans*-AD. **(A)**Monosaccharide assay of GXM from*C.n-* AstrainH99,*C.g-* B strain WM179, *C.n-* AD strain WM628, *C.g-* C strain NIH312 and *C.n-* D strain WM629 by GC-MS after uronic acid reduction treatment. **(B)** Predicted structures of GXM from five serotypes of Cryptococcus strains. **(C)** Analysis of monosaccharide composition of GXM. **(D)** ELISA assay for binding of human Control-Fc, Dectin-2-Fc, Dectin-3-Fc and Mincle-Fc fusion protein to GXM from five serotypes of Cryptococcus strains. GXM-B refers to ATCC32609 and GXM-B^#^ refers to WM179. **(E)** ELISA assay for binding of human Control-Fc and Dectin-3-Fc fusion protein to GXM from *C.g-* B strain ATCC32609(GXM-B) and *C.n-* AD strain WM628 (GXM-AD), which were performed with deacetylation, weak alkali hydrolysis and carboxyl reduction. Data are means ± SD of triplicate wells and are representative of three independent experiments; **p<0.01,***p<0.001.

To determine whether Dectin-3 directly recognizes GXM from *C.g*-B and *C.n*-AD, we used an enzyme-linked immune-sorbent assay (ELISA)-based method to examine the direct binding of plate-coated GXM extracted from *Cryptococcus* strains with Fc-fusion Dectin-2, Dectin-3 and Mincle, which were known to be bond with plate-coated α-1,2-mannan or TDM (**Figure 3D**). We found that only Dectin-3 could bind the GXM from *C.n*-AD and *C.g-* B and no binding was observed to Dectin-2 and Mincle with GXM from *C.n*-A, *C.g*-B, *C.g*-C, *C.n*-D and *C.n*-AD (**Figure 3D**). In contract, both deacetylation and weak alkali hydrolysis of GXM from *C.n*-AD and *C.g-* B, which mainly affected xylose residue and were confirmed to by ^1^H NMR and GC-MS analysis respectively (**Figure S2D** and **S2E**), significantly impaired their binding to Dectin-3 (**Figure 3E**). However, carboxyl reduction of GXM from *C.n*-AD and *C.g-* B had no any influences on their binding to Dectin-3 (**Figure 3E**). Together, these data indicate that Dectin-3 can directly recognize GXM from *C.n*-AD and *C.g-* B and xylose residue in GXM from *C.n*-AD and *C.g-* B determines their specific binding with Dectin-3.

### Dectin-3 recognizes GXM from *C.g*-B and *C.n*-AD to induce NF-κB- and ERK-mediated pro-inflammation responses

To further determine whether Dectin-3 recognizes GXM from *C.g*-B and *C.n*-AD to elicit pro-inflammation responses, we stimulated wild-type and Dectin-3-deficent BMDMs with plate-coated GXM extracted from *C.g-* B (GXM-B) and *C.n*-AD(GXM-AD). We found that stimulation with plate-coated GXM-B and GXM-AD induced IκBα phosphorylation and degradation and ERK phosphorylation (**Figure 4A and 4B**). In contract, Dectin-3 deficiency in BMDMs strikingly impaired GXM-B or GXM-AD induced activation of NF-κB and ERK pathway (**Figure 4A and 4B**). Furthermore, we found that only stimulation with plate-coated GXM-B and GXM-AD potently induced the secretion of pro-inflammation cytokines including TNF-α, IL-6 and IL-12p40 whereas Dectin-3 deficiency significantly impaired these responses (**Figure 4C**). We further examined the effect of inhibiting NF-κB or ERK signaling on pro-inflammation cytokine production by wild-type BMDMs. We found that treatment with TPCA, a specific inhibitor for NF-κB activation, significantly suppressed TNFαand IL-6 production by wild-type BMDMs when stimulated with plate-coated GXM extracted from either *C.g-* B or *C.n*-AD (**Figure 4E and 4F**). However, treatment with U0126, an inhibitor for ERK activation, only blocked TNFαproduction and had no inhibitory effect on GXM-induced IL-6 production (**Figure 4E and 4F**). Thus, these data suggest that Dectin-3 recognizes GXM from *C.g*-B and *C.n*-AD to trigger NF-κB-and ERK-mediated pro-inflammationresponses.

**Figure 4.**
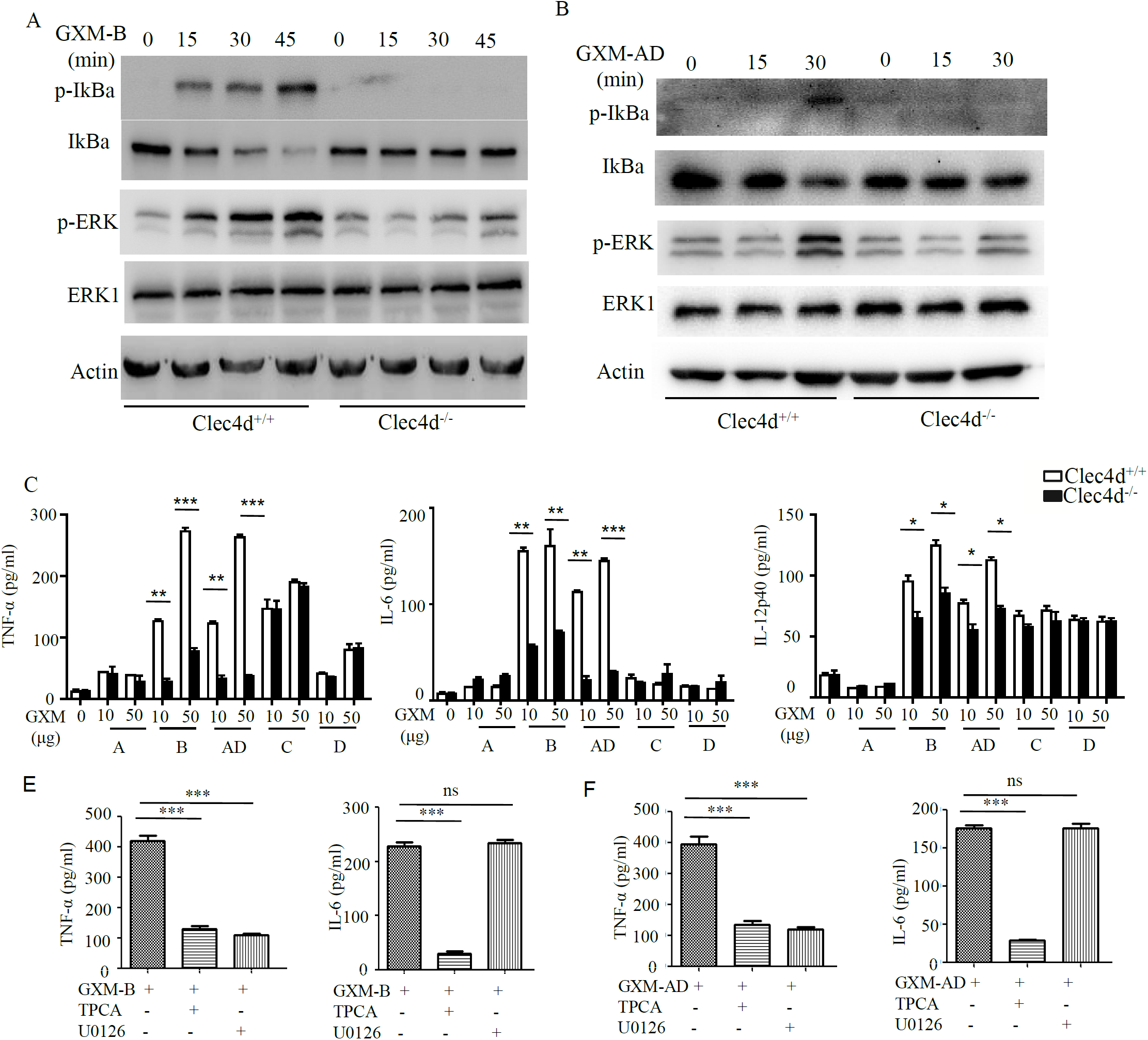
GXMs from *C.gattii*-B and *C.neoformans*-AD induce NF-κB- and ERK-mediated pro-inflammation responses through Dectin-3. **(A and B)** Protein phosphorylation levels in BMDMs from WT and Dectin-3 (Clec4d)-deficient mice, which were stimulated with plate-coated 50μg/well GXM from *C.g-* B strain ATCC32609 (GXM-B, **A**) or *C.n-* AD strain WM628 (GXM-AD, **B**) for indicated time. **(C)** ELISA results for indicated cytokines in WT and Dectin-3-deficient BMDMs, which were stimulated with plate-coated GXM extracted from *C.n*-A strain H99, *C.g-* B strain ATCC32609, *C.n-* AD strain WM628, *C.g-* C strain NIH312 and *C.n-* D strain WM629 for 16h. **(E and F)** ELISA results for indicated cytokines in WT BMDMs, which were pretreated with 10μM TPCA (p65 inhibitor) or 10μM U0126 (ERK inhibitor) and then stimulated with plate-coated GXM-B (**E**) or GXM-AD (**F**) for 16h. Data are means ± SD of triplicate wells and are representative of three independent experiments; *p<0.05, **p<0.01,***p<0.001.

### Dectin-3-deficient mice are highly susceptible to pulmonary infection with *C.g*-B and *C.n*-AD

To provide genetic evidence that Dectin-3 plays a critical role against experimental pulmonary cryptococcosis, we explored the overall impact of Dectin-3 deficiency during an experimental pulmonary infection with *C.g*-B and *C.n*-AD in mice. WT and Dectin-3 deficient mice received an intratracheal inoculation with *C.g*-B strain ATCC32609. Survival was monitored for greater than 60 days post-inoculation, while pulmonary fungal burden was evaluated in a separate group of infected mice at select time points post-inoculation (**Figure 5A and 5B**). We found that all infected Dectin-3 deficient mice died during 45 days, whereas about 70% infected WT mice survived for more than 65 days (*p* < 0.01, **Figure 5A**). Consistently, the burdens of *C.g*-B in lung and brain on day 3 after infection were significantly higher in Dectin-3 deficient mice than in WT mice(*p* <0.01 and*p* <0.001 respectively, **Figure 5B**). We then conducted a histological analysis of the lungs on day 14 after infection. Massive multiplication of yeast cells with poor granulomatous responses was observed in the alveolar spaces of Dectin-3 deficient mice, whereas WT mice showed a seldom number of yeast cells, which were mostly encapsulated in the granulomatous tissues (**Figure 5C**). We also detected the significant reductions of pro-inflammatory cytokine including TNF-α and IL-6 in the lungs of Dectin-3 deficient mice than those in WT mice (**Figure 5D**). However, the amount of IL-12p40 and IL-1β was not significantly decreased in the lungs of Dectin-3 deficient mice (**Figure 5D**). These results indicate that Dectin-3-mediated signaling is involved in the elimination of pulmonary *C.g*-B infection and induction of a protective inflammatory response.

**Figure 5.**
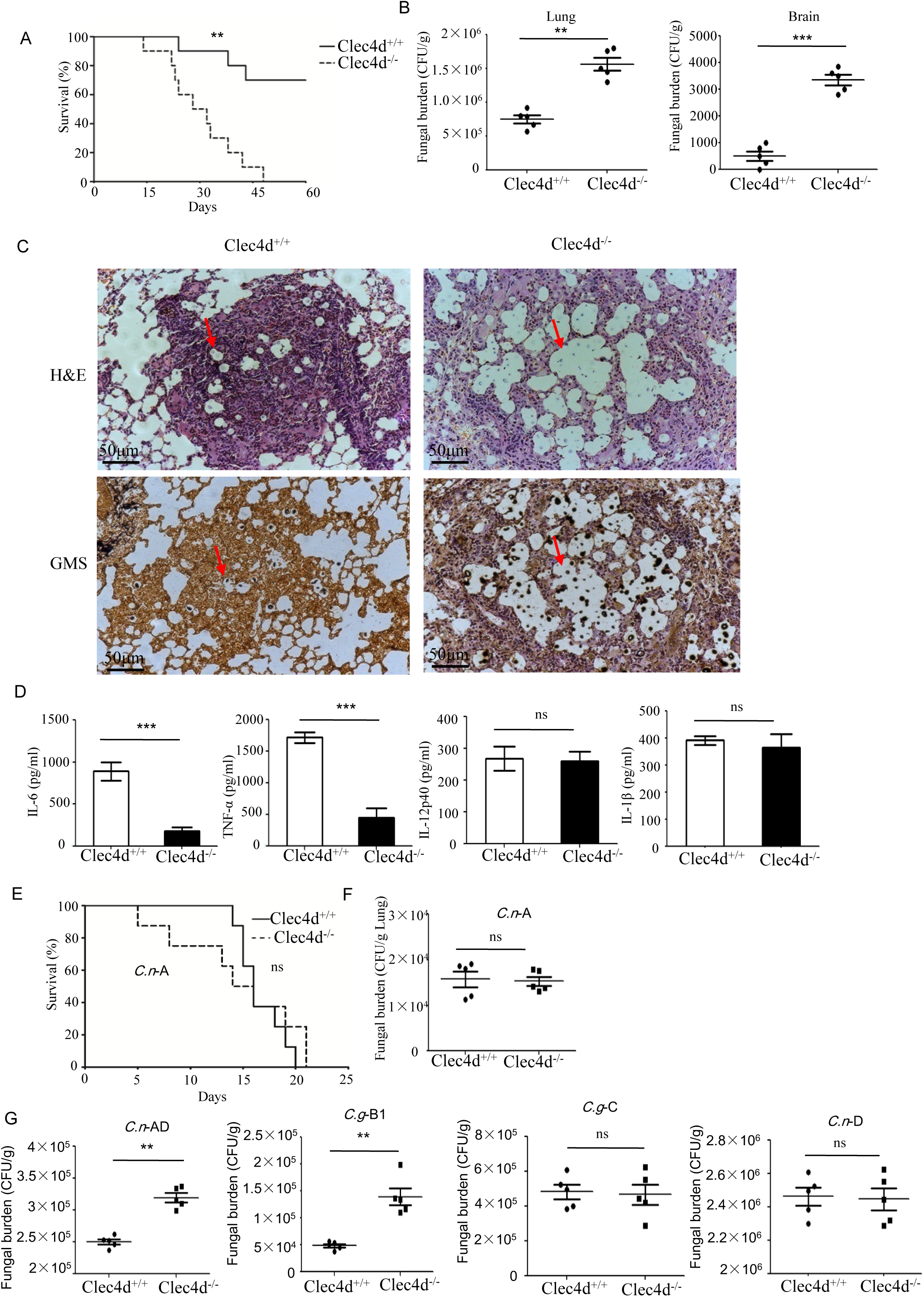
Dectin-3-deficient mice are highly susceptible to pulmonary infection with *C. gattii*-B and *C. neoformans*-AD. **(A)** Survival curves of WT and Dectin-3-deficient (Clec4d) mice (n=10 for each group) after intratracheal infection with 1×10^6^ CFU of *C.g-* B strain ATCC32609. **(B)** CFU assays of lung and brain ofWT and Dectin-3-deficient mice infected intratracheally with 1×10^5^ CFU of *C.g-* B on day 3 after infection. **(C)** Histopathology was analyzed with hematoxylin and eosin (H&E) and gomori’smethenamine silver (GMS) staining on day 14 after intratracheal infection with 1×10^6^ CFU of *C.g-* B. **(D)** ELISA results of TNF-a, IL-6, IL-12p40 and IL-1β in WT and Dectin-3-deficient mice lung homogenate on day 1 after intratracheal infection with 1×10^5^ CFU of *C.g-* B. **(E)** Survival curves of WT and Dectin-3-deficient mice (n=10 for each group) after intratracheal infection with 1×10^6^ CFU of *C.n-* A strain H99. **(F)** Lung CFU assay of WT and Dectin-3-deficient mice on day 3 after infection with 1×10^5^ CFU of *C.n-* A. **(G)** Lung CFU assays of WT and Dectin-3-deficient mice on day 3 after intratracheal infection with 1×10^5^ CFU of *C.n-* AD strain WM628, *C.g-* B strain WM179, *C.g-* C strain NIH312 or *C.n-* D strain WM629, respectively. Data are means ± SD of triplicate wells and are representative of three independent experiments; **p<0.01, ***p<0.001

Consistent with the results in the previous study [25], WT and Dectin-3 deficient mice showed an equivalent susceptibility to pulmonary *C.n*-A strain H99 infection (**Figure 5E**). Also, no significant differences were observed in the pulmonary fungal burdens of Dectin-3 deficient mice compared to WT mice on day 3 after infection with *C.n*-A strain H99 (**Figure 5F**). These results indicate that Dectin-3 is not required for host defense against pulmonary *C.n*-A infection. Moreover, Dectin-3 deficient mice exhibited higher fungal burdens in lungs than WT mice after intratracheal infection with *C.n-* AD strain WM628 and *C.g-* B strain WM179 (*p* < 0.01, **Figure 5G**). However, there were no differences in lung fungal burdens between WT and Dectin-3 deficient mice when infected with *C.g-* C strain NIH312 and *C.n-* D strain WM629 (**Figure 5G**). Together, these results demonstrate that Dectin-3-deficient mice are highly susceptible to pulmonary *C.g*-B and *C.n-* AD infections.

### Dectin-3 is critical for activation of alveolar macrophages after pulmonary *C.g*-B and *C.n*-AD infection

It has been well-documented that alveolar macrophages (AM) constitute the first line of host defense against pulmonary *Cryptococcus* infections and the subsequent inflammatory response, resulting in an influx of neutrophils and monocytes, affords a second line of defense[41]. To explore whether Dectin-3 is required for AM and neutrophil accumulation at the early stage of pulmonary infection with *C.g*-B and *C.n*-AD, We determined cellular composition of lungs and found that infected WT mice with *C.g*-B and *C.n*-AD on day 1 showed a prominent AM-biased response while Dectin-3 deficiency significantly impaired AM accumulation in lungs (*p* < 0.001, **Figure 6A and Figure S3A**). However, either pulmonary *Cryptococcus* infection or Dectin-3 deficiency had no any influences on neutrophil accumulation in lungs at the early stage (**Figure 6B and Figure S3B**). Furthermore, we found that AMs sorted from naïve WT mice had high killing activities against *C.n*-A, *C.n*-AD and *C.g*-B in a dose-dependent manner, whereas Dectin-3 deficiency significantly reduced killing activities of AMs against *C.n*-AD and *C.g*-B, but not *C.n*-A (**Figure 6C**). These data suggest that Dectin-3 controls AM accumulation and activities after pulmonary infection with *C.g*-B and*C.n*-AD.

**Figure 6.**
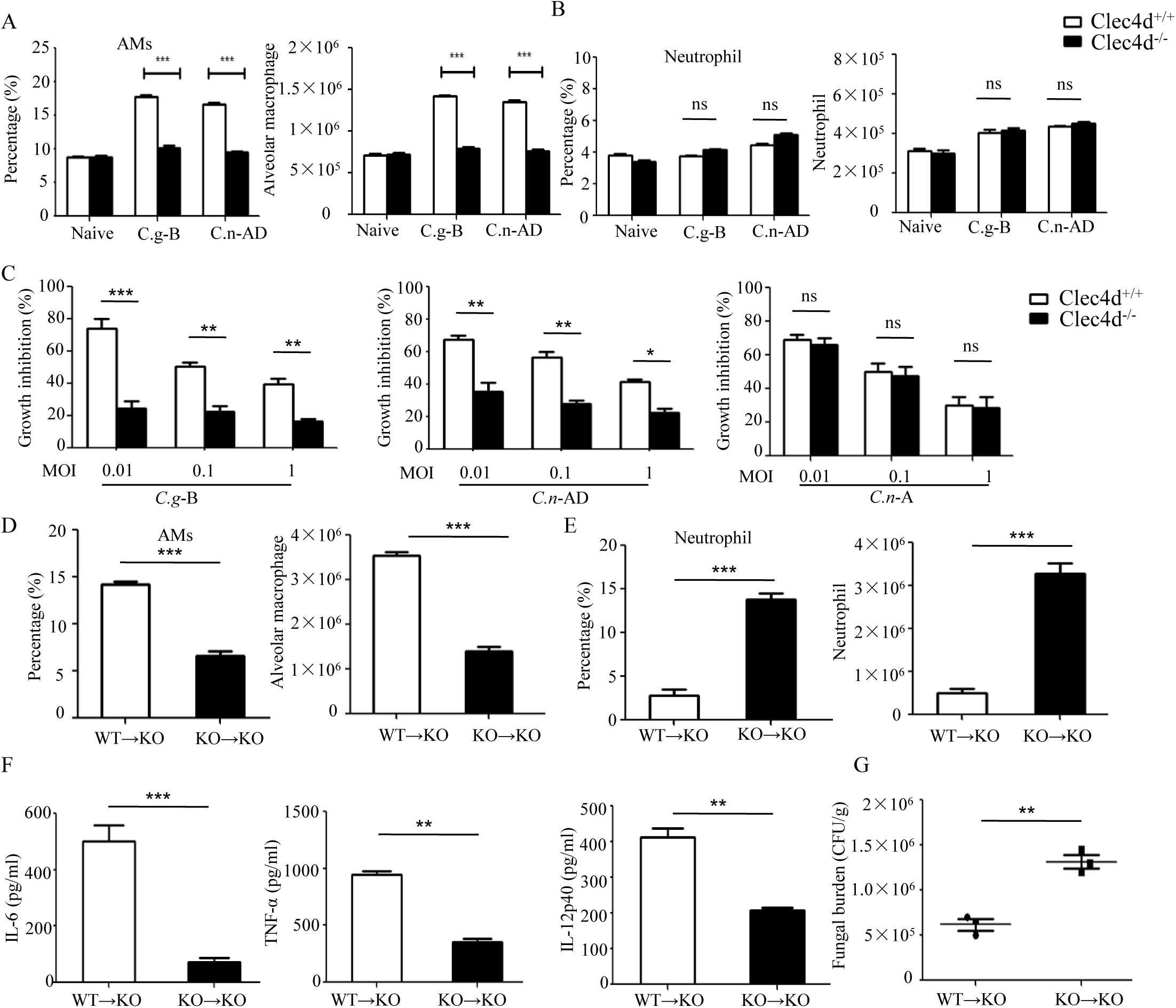
Dectin-3 is critical for activation of alveolar macrophages after pulmonary *C.gattii*-B and *C.neoformans*-AD infection. **(A and B)** Alveolar macrophages (CD11c^+^SiglecF^+^, **A**) and neutrophil (CD11b^+^Ly6G^+^, **B**) counts in lungs of WT and Dectin-3-deficient mice on day 1 after intratracheal infection with *C.g*-B strain ATCC32609 or *C.n*-AD strain WM628. **(C)** Killing activity of alveolar macrophages sorted from lungs of WT and Dectin-3-deficient mice. *C.g*-B strain ATCC32609, *C.n*-AD strain WM628 or *C.n*-A strain H99 (MOI=0.01, 0.1, 1) was co-cultured with or without sorted alveolar macrophages for 6h and the growth inhibition by alveolar macrophages was calculated as described in material and method section. **(D and E)** Alveolar macrophages (CD11c^+^SiglecF^+^, **D**) and neutrophil (CD11b^+^Ly6G^+^, **E**) counts in lungs of Dectin-3-deficient mice, which received intravenous adopt transfer of AMs (5×10^5^ cells/mouse) sorted from WT or Dectin-3-deficient mice and then infected with 1×10^5^ CFU of *C.g-* B strain ATCC32609. **(F)** ELISA results of TNF-a, IL-6 and IL-12p40 in lung homogenate of Dectin-3-deficient mice, receiving intravenous adopt transfer of alveolar macrophages, on day 1 post infection. **(G)** Lung CFU assays of Dectin-3-deficientmice, receiving intravenous adopt transfer of alveolar macrophages, on day 3 post infection.*p<0.05, **p<0.01, ***p<0.001

To further examine roles of Dectin-3 in AM-mediated host defense against pulmonary infection with *C.g*-B, we performed an adoptive transferring of equal numbers of AMs between WT and Dectin-3-deficient (KO) mice (**Figure S3C**). We found that transferred AMs sorted from WT mice into KO mice could significantly increase AM accumulation and protein levels of pro-inflammation cytokine including TNF-α, IL-6 and IL-12p40 in lung on day 1 after pulmonary infection with *C.g*-B (**Figure 6D and 6F**). Furthermore, the transfer of WT AMs into KO mice significantly reduced fungal burdens in lungs after intratracheal infection with *C.g*-B (*p* < 0.01, **Figure 6G**). In contrast, transferred AMs sorted from KO mice into KO mice had no any influences on AM accumulation, but significantly increased neutrophil accumulation in lungs (**Figure 6D and 6E**). These data confirm that Dectin-3 is critical for accumulation and activities of AM against pulmonary *C.g*-B infection.

### CARD9 is critical for pro-inflammation responses induced by GXMs

It has been shown that CARD9 operates downstream of CLRs for activating NF-κB and ERK pathways [26,27]. To explore whether CARD9 is required for*C.g*-B-induced pro-inflammation responses, we stimulated BMDMs from WT and CARD9 deficient mice with plate-coated GXM extracted from *C.g-* B (GXM-B), and found that CARD9 deficiency in BMDMs completely impaired GXM-B induced activation of NF-κB and ERK pathway (**Figure S4A and S4B**). Furthermore, we found that CARD9 deficiency in BMDMs significantly impaired GXM-B induced secretion of pro-inflammation cytokines including TNF-α and IL-6, but not IL12-p40 and IL-1β(**Figure 7A**). Moreover, CARD9 deficiency in BMDMs also significantly impaired the production of TNF-α and IL-6 when stimulated with plate-coated GXM extracted from *C.n*-AD, *C.g*-C and *C.n*-D, but not *C.n*-A (**Figure 7A**). These data suggest that CARD9 is essential for NF-κB and ERK-mediated pro-inflammation responses induced by GXM from *C.g*-B, *C.g*-C, *C.n*-D and *C.n*-AD.

**Figure 7.**
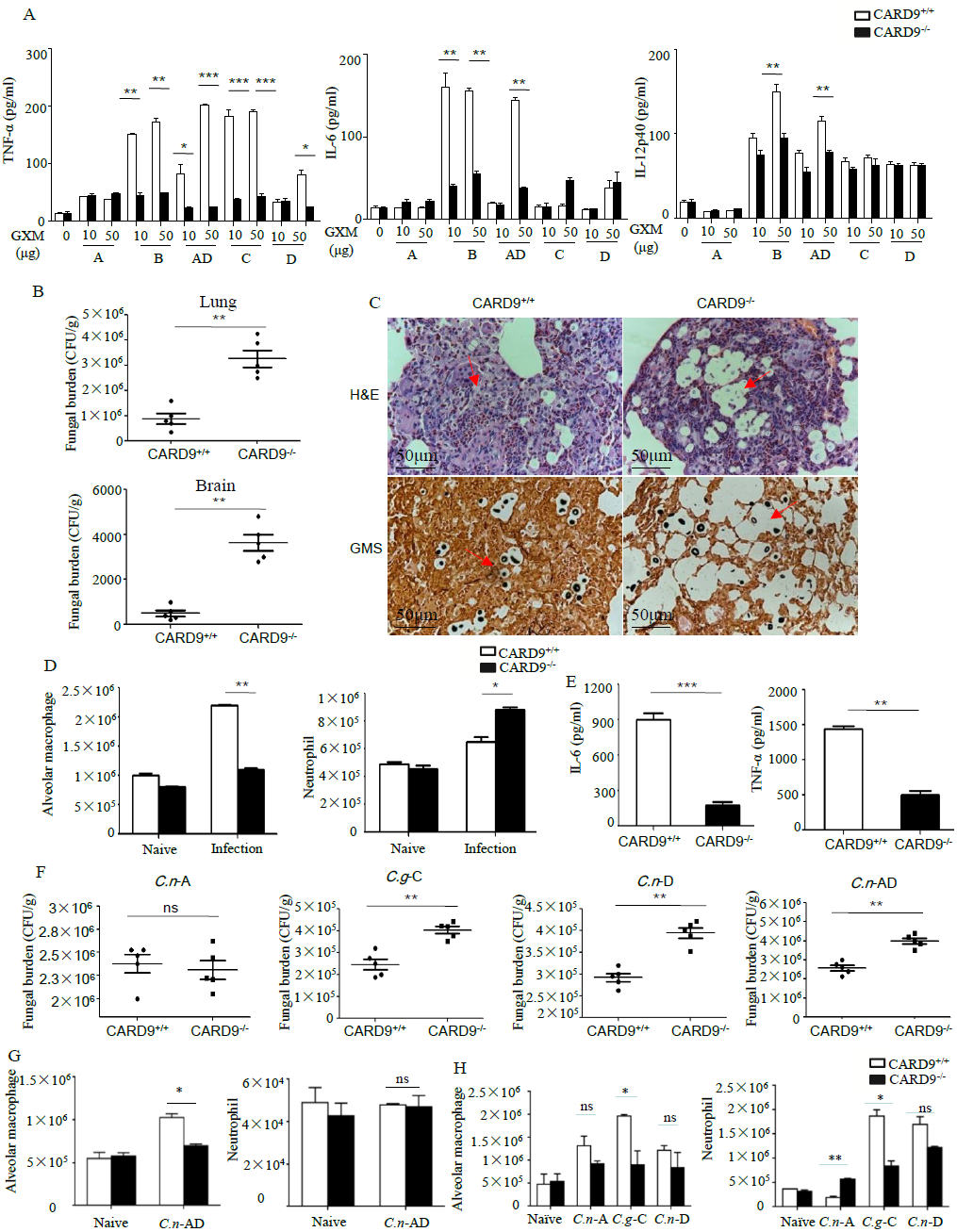
CARD9 is critical for pro-inflammation responses induced by GXMs. **(A)** ELISA results for indicated cytokines in WT and CARD9-deficient BMDMs, which were stimulated with plate-coated GXM extracted from *C.n-* A strain H99, *C.g-* B strain ATCC32609, *C.n-* AD strain WM628, *C.g-* C strain NIH312 and *C.n-* D strain WM629 for 16h. **(B)** CFU assays of lung and brain of WT and CARD9-deficient mice after intratracheal infection with 1×10^5^ CFU of *C.g-* B strain ATCC32609 on day 3 post infection. **(C)** Histopathology was analyzed with H&E and GMS staining on day 14 after intratracheal infection with 1×10^6^ CFU of *C.g-* B strain ATCC32609. **(D)** Alveolar macrophages (CD11c^+^SiglecF^+^) and neutrophil (CD11b^+^Ly6G^+^) counts in lungs of WT and CARD9-deficient mice on day 1 after intratracheal infection with *C.g*-B strain ATCC32609. **(E)** ELISA results of TNF-a and IL-6 in lung homogenate of WT and CARD9-deficient mice on day 1 after infection. **(F)** Lung CFU assays of WT and CARD9-deficient mice on day 3 after intratracheal infection with 1×10^5^ CFU of *C.n-* AD strain WM628, *C.g-* B strain WM179, *C.g-* C strain NIH312 or *C.n-* D strain WM629, respectively. **(G and H)** Alveolar macrophages (CD11c^+^SiglecF^+^) and neutrophil (CD11b^+^Ly6G^+^) counts in lungs of WT and CARD9-deficient mice on day 1 after intratracheal infection with *C.n-* AD strain WM628 **(G)** or *C.n-* A strain H99, *C.g-* C strain NIH312 and *C.n-* D strain WM629 **(H)**. Data are means ± SD of triplicate wells and are representative of three independent experiments; *p<0.05, **p<0.01, ***p<0.001

To examine the effect of CARD9 deficiency on the clinical course of cryptococcal infection, we intratracheally infected WT and CARD9 deficient mice with different *Cryptococcus* strains, and found that the burdens in lung and brain were significantly higher in CARD9 deficient mice than those in WT mice on day 3 and 14 after infection with *C.g*-B strain ATCC32609(**Figure 7B and Figure S4C**). Furthermore, the histological analysis of the lungs on day 14 after infection showed massive multiplication of yeast cells with poor granulomatous responses in the alveolar spaces of CARD9 deficient mice (**Figure 7C**), which is consistent with the findings observed in infected Dectin-3 deficient mice with *C.g*-B strain ATCC32609 (**Figure 5C**). Moreover, CARD9 deficiency completely blocked AM accumulation (**Figure 7D and Figure S4D**) and significantly reduced secretion of pro-inflammatory cytokine including TNF-α and IL-6 in the lungs on day 1 after infection with *C.g*-B (**Figure 7E**). Unexpectedly, CARD9 deficiency slightly increased neutrophil accumulation in the lungs (**Figure 7D and Figure S4E**), which has been reported in a previous study [28]. Moreover, CARD9 deficient mice exhibited higher fungal burdens in lungs than WT mice after intratracheal infection with *C.n-* AD strain WM628, *C.g-* C strain NIH312 and *C.n-* D strain WM629 for 3 days (*p* < 0.01, **Figure 7F**). However, there were no differences in lung fungal burdens between WT and CARD9 deficient mice when infected with *C.n-* A strain H99 on day3(**Figure7F**). Moreover, CARD9 deficiency completely blocked AM accumulation in the lungs at day 1 after infection with *C.n-* AD strain WM628 and *C.g-* C strain NIH312 (**Figure 7G and 7H, Figure S5 and S6**). Unexpectedly, CARD9 deficiency had no influences on AM accumulation in the lungs at day 1 after infection with *C.n-* D strain WM629 (**Figure 7H and Figure S6**). These data show that CARD9 is critical for accumualtion and activities of AM against pulmonary infection with *C.g*-B, *C.g-* Cand *C.n-* AD.

## Discussion

Germline encoded PRRs recognize a variety of microbial moieties which, once engaged, result in the activation of anti-microbial host defense and stimulation of adaptive immune responses. The role of CLRs during cryptococcosis are of interest as recent studies have defined their role in the recognition of carbohydrate moieties and host defense against other fungal pathogens[15,16,18,21,23,42]. In the present study, we showed that Dectin-3-deficient mice were highly susceptible to pulmonary infection with *C. g-* B or *C. n-* AD strains as shown by the increased fungal burdens in lung and brain. These mice also showed massive multiplication of yeast cells in the alveolar spaces with poor granulomatous responses and the significant reductions of pro-inflammatory cytokine including TNF-α and IL-6 in the lungs. Moreover, Dectin-3 deficiency significantly impaired accumulation and killing activity of alveolar macrophages (AM) in lungs, whereas adoptive transferring of WT AM into Dectin-3 deficient mice conferred their resistance to pulmonary infection with *C. g-* B. Thus, Dectin-3-mediated signaling was demonstrated to be critical for rendering the host defense against pulmonary infection with *C.g*-B and *C.n*-AD. Nonetheless, we observed no defect in the capacity of Dectin-3-deficient mice to combat experimental pulmonary infections with *C. n-* A, *C. n-* D or *C. g-* C, which is in line with the previous study [25]. Thus, our results suggest that the role of Dectin-3 for host defense against cryptococcosis is specific to *Cryptococcus* serotypes. However, Dectin-1, which recognizes β-1,3-glucans in a variety of fungi including *C*. *albicans* [15], is not required for protection against pulmonary *C. n-* A infection[17]. Dectin-2 is postulated to induce Th2-type responses and IL-4-dependent mucin production in the lungs following infection with *C. n-* D[20]. The impact of Dectin-1 or Dectin-2 deficiency may also vary depending on the *Cryptococcus* serotypes.

*Cryptococcus* strains possess the large polysaccharide capsule to shield its cell wall components and the capsule is primarily composed of GXM, which comprises more than 90% of the capsular polysaccharide mass[29]. The cell wall of *Cryptococcus* strains is composed of chitin, chitosan, glucans and glycoproteins. Thus, carbohydrates present in the capsule and cell wall of *Cryptococcus* strains are ideal PAMPs for recognition by Dectin-3. Our present study showed that Dectin-3 could directly recognize the capsular GXM extracted from *C. g-* B or *C. n-* AD strains. Monosaccharide composition analysis showed the quantity and position of xylose residue was major structural difference of GXMs among *C.n*-A, *C.g*-B, *C.g*-C, *C.n*-D and *C.n*-AD. In detail, GXMs from *Cryptococcus* strains are all comprised of a core repeating unit to which (1→2)-linked and (1→4)-linked β-D-Xyl*p* units are added in increments of one to four residues. And GXMs from *C.n*-A and *C.n*-D are mainly substituted with Xylp at C-2, whereas GXMs from *C.g*-B, *C.g*-C and *C.n*-AD are substituted at C-2 and at C-4. Thus, these structural differences may determine that Dectin-3 differentially recognizes GXM from *C.n*-A, *C.g*-B, *C.g*-C, *C.n*-D and *C.n*-AD.

Furthermore, we showed that Dectin-3 mediated GXM-induced inflammatory responses through activating NF-κB and ERK pathways. And inhibiting NF-κB or ERK signaling could strikingly block pro-inflammatory responses induced by GXM extracted from *C. g-* B or *C. n-* AD strains. Our previous studies show that Dectin-3 stimulation with α1,2-mannans of *C. albicans* hyphae or by TDM of *M. tuberculosis* could facilitate activation of NF-κB pathway[23,24]. However, it has not been characterized how Dectin-3 signaling pathway leads to ERK activation. Our previous study also show that stimulation of Dectin-1 with β1,3-glucans of *C. albicans* yeast induces CARD9-mediated ERK activation by linking Ras-GRF1 to H-Ras[27]. Our present study showed that CARD9 deficiency dramatically impaired activation of both NF-κB and ERK pathway induced by GXM extracted from *C. g-* B strain and subsequently blocked pro-inflammation cytokine production induced by GXM from *C.g*-B, *C.g*-C, *C.n*-D and *C.n*-AD. Furthermore, our data showed that CARD9 deficient mice were highly susceptible to pulmonary infection with *C.g*-B, *C.g*-C, *C.n*-D and *C.n*-AD, as shown by the increased fungal burdens in the infected lungs at the early stage. CARD9 deficient mice infected with *C.g*-B showed remarkable multiplication of the yeast cells in the alveolar spaces with poor granulomatous responses. Moreover, CARD9 deficiency significantly impaired AM accumulation and the production of pro-inflammatory cytokine including TNF-α and IL-6 in the lungs. Thus, CARD9-mediated signaling was also critical for rendering the host defense against pulmonary infection with *C.g*-B, *C.g*-C, *C.n*-D and *C.n*-AD. However, PRRs for recognizing *C.g*-C and *C.n*-D remain to be explored in the future.

In conclusion, we have extensively characterized the role of Dectin-3 and CARD9 during pulmonary cryptococcosis using an experimental murine model of pulmonary infection with *C. neoformans* and *C. gattii*. Upon pulmonary *C.n*-AD and *C.g*-B infection, Dectin-3- and CARD9-deficient mice were highly susceptible and showed augmented lung injury due to impairment of AM accumulation and killing activities, and subsequent productions of pro-inflammation cytokines and chemokines. More importantly, we showed that Dectin-3 directly recognized GXMs from *C.n*-AD and *C.g*-B to activate CARD9-mediated NF-κB and ERK pathways for initiating anti-fungal host responses against *C. neoformans* and *C. gattii* **(Figure 8)**. Thus, our study provides the first biological and genetic evidence demonstrating that Dectin-3 recognizes GXM of *C. neoformans* serotype AD *and C. gattii* serotype B to initiate host defense against cryptococcosis.

**Figure 8.**
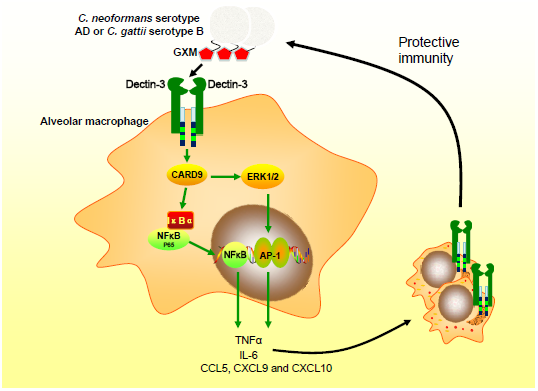
Proposed model that Dectin-3 recognized GXMs from *C.n*-AD and *C.g*-B to activate CARD9-mediated NF-κB and ERK pathways.

## Materials and Methods

### Cryptococcus strain

*C.gattii-* B strain ATCC32609 and WM179, *C.gattii-* C strain NIH312, *C.neoformans-* A strain H99, *C.neoformans-* D strain WM629 and *C.neoformans-* AD strain WM628 were gifted from Shanghai institute of fungal medicine, China. The yeast cells were cultured on Sabouraud Dextrose Agar (SDA) plates before use. Strains cultured in Yeast Extract Peptone Dextrose (YPD) medium at 30°C for 16h, were presented as thin-capsule form. Strains cultured in RPMI-1640 medium plus 10% FBS at 37°C with 5% CO_2_ for 16h, were presented as thick-capsule form.

### BMDM preparation

Primary cultures of BMDMs from C57B/L6 mice were prepared as previously described [43]. Briefly, bone marrow cells were harvested from the femurs and tibias of mice. Erythrocytes were removed from cells by using a hypotonic solution. Cells were cultured for 7 days in DMEM medium containing 20%fetal bovine serum, 50 mM β-mercaptoethanol, 100μg/mL penicillin-streptomycin, and 30% conditioned medium from L929 cells expressing M-CSF. On day 3, another 10 ml of the same medium was added. On day 7, non-adherent cells were removed, and adherent cells were harvested.

### Chemokine assay

Total RNA was extracted from the infected lungs or BMDMs using Triozl (TaKaRa Bio, Otsu, Japan), and the first-strand cDNA was synthesized using PrimeScript first-strand cDNAsynthesis kit (TaKaRa Bio, Otsu, Japan), according to the manufacturer’s instructions. Quantitative real-time PCR was performed in a volume of 20 μl using gene-specific primers and SYBR Premix Ex TaqII (TaKaRa Bio, Otsu, Japan) in a ABI 7500 system (Applied Biosystems, USA). The amounts of transcript were normalized to GAPDH. ΔΔCt method to calculate fold changes. First, Ct target gene - Ct housekeeping gene = ΔCt. Second, ΔCt treatment -ΔCt control = ΔΔCt. Third, fold changes = 2^−ΔΔCt^.

### Cytokine assay

TNF-a, IL-6, IL-12p40 and IL-1β concentrations in the lung homogenates and culture supernatants were measured by SET-Ready-GO ELISA kits (eBioscience) according to the manufacturer’s protocol.

### Immunoblot analysis

Immunoblotting was performed as described previously [27]. Anti-IκBα (9247),-phospho-IκBα (9246),-ERK (4372), -phospho-ERK (4695), - NF-κB-p65 (8242) antibodies were obtained from Cell Signaling (Danvers, MA). Anti-β-actin (ab8224), -PCNA (ab152112) antibodies were from Abcam (Cambridge, MA).

### Isolation and purification of GXM

The extracted extracellular polysaccharides (EPS) from *Cryptococcus* strains were isolated as previously described [44]. In detail, strains were grown in YNB culture medium at 30°C with shaking for 4 days and then the supernatant were collected after centrifuging with 7500 rpm for 5min. The supernatant was slowly added with 3 volumes of EtOH and incubated for overnight at 4°C, which were centrifuged at 7500 rpm for 10min. The precipitate was collected for air dry and then dissolved into 18 mL ddH_2_O. The polysaccharide concentration was determined by the phenol sulfuric method of Dubois (data not shown). To purify GXM, the polysaccharide solution was adjusted to 0.2M NaCl and finalized with 3 times (w/w) of 0.3% (w/v) CTAB solution at room temperature, the mixture of which is named as CTAB-GXM. Then, CTAB-GXM was centrifuged at 9000 rpm for 10 min and the precipitate of CTAB-GXM was washed with 10% EtOH in ddH20 (v/v). To remove CTAB, the precipitate of CTAB-GXM was dissolved in 1 M NaCl and finalized with about 2.5 times (v/v) of EtOH with stirring. The precipitate containing GXM was collected after centrifuging at 9000 rpm for 10 min at room temperature and then dissolved in 2 M NaCl until a viscous solution forms. The viscous solution was dialyzed in a dialysis cassette with a 10,000 MW cutoff for overnight against 1 M NaCl and then against fresh ddH_2_O for 1 week. Finally, the solution EPS containing GXM inside the dialysis bag was lyophilized at −55°C for overnight. Endotoxin levels of GXM were measured by endotoxin assay kit (GenScript, Cat. No.L00350C).

### Analysis of monosaccharide composition of GXM

EPS (1mg) was hydrolyzed with 2 M TFA at 120°C for 2h. After repeated evaporation with methanol to completely remove TFA, the residue was dissolved in distilled water and reduced with NaBH_4_ at room temperature for 3h. After neutralization with AcOH and evaporation to dryness, the residue was acetylated with Ac_2_O for 1h at 100°C. The resulting alditol acetates were analyzed with gas chromatography–mass spectrometry (GC-MS). The GC-MS temperature program for monosaccharide analysis was as follows: 140-198°C at 2°C/min, maintained for 4 min and increased to 214°C at 4°C /min, followed by increases of 1°C /min until 217°C was reached, which was maintained for 4 min and finally increased to 250°C at 3°C/min, maintained for 5 min as reported previously [45].

### Chemical modification of GXM

EPS was performed with 1-cyclohexyl-3-(2-morpholinoethyl) carbodiimide metho-*p*-toluenesulfonate (CMC) and NaBH4 for carboxyl reduction[46]. This procedure resulted in D-glucuronic acid converted into D-glucose. For deacylation, EPS (25mg) was treated with 5 ml NaOH (0.1M) for 16h at 37°C, neutralized with 0.1M HCl, and then dialyzed against water [47]. For partial acid hydrolysis, EPS (50 mg) was hydrolyzed in 5ml TFA (0.1M) and incubated at 100°C for 1h. The hydrolysate solution was evaporated to remove TFA through repeated evaporation with MeOH under reduced pressure [45].

### NMR analysis

EPS (80 mg) was dissolved in 0.4 ml D_2_O (99.8 Atom% Deuterium; Schweres Wasser, USA). The ^1^H nuclear magnetic resonance (NMR) spectra were measured using the Bruker AvanceIII 600 Spectrometer (Bruker Instruments, Inc., Billerica, MA, USA) at 25°C. The chemical shifts of ^1^H NMR are expressed in ppm by using acetone as an internal standard; 4.70 ppm for ^1^H NMR. All the experiments were recorded, and data were processed using standard Bruker software and MestReNova[48].

### Expression of fusion protein

The hCLRs-pFUSE-hIgG1-Fc vectors were then transfected into 293T using Lipo2000 (Sigma), one days later, adding bleomycin (invivoGen), the best proportion of bleomycin VS. DMEM is 3:1000. After culturing three days, the cell supernatant was collected and the expression of CLR-Fc fusion protein was determined by immunoblot analysis with anti-Fc antibody.

### FACS

Yeast cells underwent lag phase induction and were plated at 1×10^7^ cells/tube. Staining was performed in 0.5% BSA in PBS. Fusion proteins were then applied at 500μL/tube followed by anti-human FITC-Fc secondary antibody (Abcam, Cambridge, MA, USA). Median fluorescence intensity (MFI) of each tube was measured by BD FACSArray flow cytometer (BD Biosciences, San Jose, CA) and presented as histogram.

### ELISA binding assay

Soluble GXM was adsorbed to ELISA plate at the concentration of 10μg/ml. Then, plate was blocked with 5% BSA at room temperature for 1h. Fusion proteins were added with 100μl/well, incubated at 37°C for 2h, and followed by 100μl/well anti-human HRP-Fc secondary antibody (Jackson ImmunoResearch Laboratories, West Grove, PA, USA) at room temperature for 30min. There were three times of washing between each step. After that, 100μl/well substrates were added at room temperature for 10min. Finally, 50μl/well 1M phosphoric acid were added as stop solution. Test results were measured on a plate reader at 450nm.

### Mice

WT C57BL/6, Dectin-3-deficient (Clec4d^−/−^) and CARD9-deficient (CARD9^−/−^) mice were kept under specific-pathogen-free conditions at the Institute for Animal Experimentation, Tongji University School of Medicine. 6-8 weeks old male and female mice were used in this study. No evidence of susceptibility based on sex was observed.

### Pulmonary cryptococcal infection in mice

Mice were anesthetized by inhaling isoflurane (RWD life science, Shenzhen, China) and restrained on a small board. Live yeast cells (1×10^5^ or 1×10^6^) were inoculated in a 35μL volume into the trachea of each mouse.

### Fungal burden analysis

Mice were sacrificed on day 3 after infection, and lungs or brains were dissected carefully, excised, and then homogenized separately in 1ml of distilled water by teasing with a stainless mesh at room temperature. The homogenates, diluted appropriately with distilled water, were inoculated at 100μl on SDA plates and cultured for 2 days, and the resulting colonies were counted.

### Histological examination

The lung specimens obtained from mice were fixed in 10% buffer formalin, dehydrated, and embedded in paraffin. Sections were cut and stained with hematoxylin-eosin (H-E) or gomori’s methenamine silver (GMS) stain, using standard staining procedures, at pathology platform of Servicebio Technology, Wuhan, China.

### Flow cytometry and cell sorting

Anti-mouse antibodies to CD45-FITC, CD11b-PerCP-Cy5.5, CD11c-APC, Ly6G-BV421, Siglec-F-PE and fixable viability stain 780 were obtained from BD Pharmingen. The lungs from mice were perfused and digested into single-cell suspensions as described previously[49]. After RBC lysis buffer treatment, the whole lung cells were washed with PBS and then stained with corresponding fluorescent antibodies. Following incubation, samples were washed and fixed in 2% ultrapure formaldehyde. The absolute number of each leukocyte subset was then determined by multiplying the absolute number of CD45^+^ cells by the percentage of cells stained by fluorochrome-labeled antibodies for each cell population analyzed using BD FACSArray software(tm) on a BD FACSArray flow cytometer (BD Biosciences, San Jose, CA). Alveolar macrophages were identified as CD45^+^CD11c^+^Siglec-F^+^. Neutrophils were identified asCD45^+^CD11b^+^Ly6G^+^. Cell sorting was performed on BD FACSAria II instrument, using BD FACSDiva software (BD Biosciences), and compensation and data analyses were performed using FlowJo software (TreeStar, Ashland, OR). Cell populations were identified using sequential gating strategy (see supplementary results). The sorting purity was 90%-95%.

### *In vitro* killing assay

Alveolar macrophages (AMs) were sorted from the lung leukocytes of mice. The purified AMs (1×10^4^/well) were cultured, in triplicate, within individual wells of a 96-well U-bottom tissue culture plate in DMEM complete media containing *C.n*-A, *C.g*-B and *C.n*-AD respectively at multiplicity of infection (MOI) of 1, 0.1, 0.01; and with *C.n*-A, *C.g*-B and *C.n*-AD in DMEM complete media without AMs as a control. After 6h, the content of each well was centrifuged and the supernatants removed. The AMs in the cell pellet were then lysed by washing 3 times with sterile water and incubating in water for 20 min. The remaining yeast was diluted in PBS and plated onto SDA plate to quantify the live cryptococcal cells.

### Ethics statement

All animal experimental procedures were performed in accordance with the Regulations for the Administration of Affairs Concerning Experimental Animals approved by the State Council of People’s Republic of China. The protocol was approved by the Institutional Animal Care and Use Committee of Tongji University (Permit Number: TJLAC-015-002). Human peripheral blood mononuclear cells (PBMCs) were obtained from Department of Medical Laboratory, Shanghai Pulmonary Hospital of China.

### Statistical analysis

At least two biological replicates were performed for all experiments unless otherwise indicated. Log-rank testing was used to evaluate the equality of survival curves. Student’s t-test for paired observations was used for statistical analyses of cytokine expression levels. Statistical significance was set at a *P* value of less than 0.05, 0.01 or 0.001.

## Acknowledgments

We thank Dr. Xin Lin and Dr. Xue-Qiang Zhao for helpful advice and suggestions. This work was supported by the National Natural Science Foundation of China (31622023 and 81571611 to X.M.J), Shanghai laboratory animal research fund (16140902600 to X.M.J), Outstanding academic leaders of Shanghai health and Family Planning Commission (2017BR024 to X.M.J), and Shuguang Program of Shanghai Education Development Foundation and Shanghai Municipal Education Commission (17SG24 to X.M.J).

## Author contributions

H.-R.H., and X.-M.J. designed the experiments. H.-R.H., F.L., H.H., Q.-Z.L., X.X., N.L. and S.W. performed the experiments. H.-R.H., J.F.X. and X.-M.J. analyzed the data. H.-R.H. and X.-M.J. wrote the manuscript with editorial input from all theauthors.

## References

1. Idnurm A, Bahn Y-S, Nielsen K, Lin X, Fraser JA, et al. (2005) Deciphering the model pathogenic fungus Cryptococcus neoformans. Nature reviews Microbiology 3: 753.

2. Kwon-Chung KJ, Boekhout T, Fell JW, Diaz M (2002) (1557) Proposal to conserve the name Cryptococcus gattii against C. hondurianus and C. bacillisporus (Basidiomycota, Hymenomycetes, Tremellomycetidae). Taxon 51: 804–806.

3. Dromer F, Gueho E, Ronin O, Dupont B (1993) Serotyping of Cryptococcus neoformans by using a monoclonal antibody specific for capsular polysaccharide. Journal of clinical microbiology 31: 359–363.

4. Ikeda R, Nishimura S, Nishikawa A, Shinoda T (1996) Production of agglutinating monoclonal antibody against antigen 8 specific for Cryptococcus neoformans serotype D. Clinical and diagnostic laboratory immunology 3: 89–92.

5. Mitchell TG, Perfect JR (1995) Cryptococcosis in the era of AIDS-100 years after the discovery of Cryptococcus neoformans. Clinical microbiology reviews 8: 515–548.

6. Dromer F, Mathoulin-Pélissier S, Launay O, Lortholary O, Group FCS (2007) Determinants of disease presentation and outcome during cryptococcosis: the CryptoA/D study. PLoS medicine 4: e21.

7. Viviani MA, Cogliati M, Esposto MC, Lemmer K, Tintelnot K, et al. (2006) Molecular analysis of 311 Cryptococcus neoformans isolates from a 30-month ECMM survey of cryptococcosis in Europe. FEMS yeast research 6: 614–619.

8. Galanis E, Hoang L, Kibsey P, Morshed M, Phillips P (2009) Clinical presentation, diagnosis and management of Cryptococcus gattii cases: Lessons learned from British Columbia. Can J Infect Dis Med Microbiol 20: 23–28.

9. Kwon-Chung K, Bennett J (1984) Epidemiologic differences between the two varieties of Cryptococcus neoformans. American journal of epidemiology 120: 123–130.

10. Kidd S, Hagen F, Tscharke R, Huynh M, Bartlett K, et al. (2004) A rare genotype of Cryptococcus gattii caused the cryptococcosis outbreak on Vancouver Island (British Columbia, Canada). Proceedings of the National Academy of Sciences of the United States of America 101: 17258–17263.

11. Byrnes III EJ, Li W, Lewit Y, Ma H, Voelz K, et al. (2010) Emergence and pathogenicity of highly virulent Cryptococcus gattii genotypes in the northwest United States. PLoS pathogens 6: e1000850.

12. Goldman D, Khine H, Abadi J, Lindenberg D, Pirofski L, et al. (2001) Serologic evidence for Cryptococcus neoformans infection in early childhood. Pediatrics 107: E66.

13. Hardison SE, Brown GD (2012) C-type lectin receptors orchestrate antifungal immunity. Nat Immunol 13: 817–822.

14. Geijtenbeek TB, Gringhuis SI (2016) C-type lectin receptors in the control of T helper cell differentiation. Nat Rev Immunol 16: 433–448.

15. Taylor PR, Tsoni SV, Willment JA, Dennehy KM, Rosas M, et al. (2007) Dectin-1 is required for beta-glucan recognition and control of fungal infection. Nat Immunol 8: 31–38.

16. Steele C, Rapaka RR, Metz A, Pop SM, Williams DL, et al. (2005) The beta-glucan receptor dectin-1 recognizes specific morphologies of Aspergillus fumigatus. PLoS Pathog 1: e42.

17. Nakamura K, Kinjo T, Saijo S, Miyazato A, Adachi Y, et al. (2007) Dectin-1 is not required for the host defense to Cryptococcus neoformans. Microbiol Immunol 51: 1115–1119.

18. Saijo S, Ikeda S, Yamabe K, Kakuta S, Ishigame H, et al. (2010) Dectin-2 recognition of alpha-mannans and induction of Th17 cell differentiation is essential for host defense against Candida albicans. Immunity 32: 681–691.

19. Loures FV, Rohm M, Lee CK, Santos E, Wang JP, et al. (2015) Recognition of Aspergillus fumigatus hyphae by human plasmacytoid dendritic cells is mediated by dectin-2 and results in formation of extracellular traps. PLoS Pathog 11: e1004643.

20. Nakamura Y, Sato K, Yamamoto H, Matsumura K, Matsumoto I, et al. (2015) Dectin-2 deficiency promotes Th2 response and mucin production in the lungs after pulmonary infection with Cryptococcus neoformans. Infect Immun 83: 671–681.

21. Ishikawa T, Itoh F, Yoshida S, Saijo S, Matsuzawa T, et al. (2013) Identification of distinct ligands for the C-type lectin receptors Mincle and Dectin-2 in the pathogenic fungus Malassezia. Cell Host Microbe 13: 477–488.

22. Yonekawa A, Saijo S, Hoshino Y, Miyake Y, Ishikawa E, et al. (2014) Dectin-2 is a direct receptor for mannose-capped lipoarabinomannan of mycobacteria. Immunity 41: 402–413.

23. Zhu LL, Zhao XQ, Jiang C, You Y, Chen XP, et al. (2013) C-type lectin receptors Dectin-3 and Dectin-2 form a heterodimeric pattern-recognition receptor for host defense against fungal infection. Immunity 39: 324–334.

24. Zhao XQ, Zhu LL, Chang Q, Jiang C, You Y, et al. (2014) C-type lectin receptor dectin-3 mediates trehalose 6,6’-dimycolate (TDM)-induced Mincle expression through CARD9/Bcl10/MALT1-dependent nuclear factor (NF)-kappaB activation. J Biol Chem 289: 30052–30062.

25. Campuzano A, Castro-Lopez N, Wozniak KL, Leopold Wager CM, Wormley FL Jr,. (2017) Dectin-3 Is Not Required for Protection against Cryptococcus neoformans Infection. PLoS One 12: e0169347.

26. Gross O, Gewies A, Finger K, Schafer M, Sparwasser T, et al. (2006) Card9 controls a non-TLR signalling pathway for innate anti-fungal immunity. Nature 442: 651–656.

27. Jia XM, Tang B, Zhu LL, Liu YH, Zhao XQ, et al. (2014) CARD9 mediates Dectin-1-induced ERK activation by linking Ras-GRF1 to H-Ras for antifungal immunity. J Exp Med 211: 2307–2321.

28. Yamamoto H, Nakamura Y, Sato K, Takahashi Y, Nomura T, et al. (2014) Defect of CARD9 leads to impaired accumulation of gamma interferon-producing memory phenotype T cells in lungs and increased susceptibility to pulmonary infection with Cryptococcus neoformans. Infect Immun 82: 1606–1615.

29. Zaragoza O, Rodrigues ML, De Jesus M, Frases S, Dadachova E, et al. (2009) The capsule of the fungal pathogen Cryptococcus neoformans. Adv Appl Microbiol 68: 133–216.

30. Doering TL (2000) How does Cryptococcus get its coat? Trends in microbiology 8: 547–553.

31. Bhattacharjee AK, Bennett JE, Glaudemans CP (1984) Capsular polysaccharides of Cryptococcus neoformans. Rev Infect Dis 6: 619–624.

32. Cherniak R, Valafar H, Morris LC, Valafar F (1998) Cryptococcus neoformans chemotyping by quantitative analysis of 1H nuclear magnetic resonance spectra of glucuronoxylomannans with a computer-simulated artificial neural network. Clin Diagn Lab Immunol 5: 146–159.

33. Belay T, Cherniak R (1995) Determination of antigen binding specificities of Cryptococcus neoformans factor sera by enzyme-linked immunosorbent assay. Infect Immun 63: 1810–1819.

34. Shoham S, Huang C, Chen JM, Golenbock DT, Levitz SM (2001) Toll-like receptor 4 mediates intracellular signaling without TNF-alpha release in response to Cryptococcus neoformans polysaccharide capsule. J Immunol 166: 4620–4626.

35. Yauch LE, Mansour MK, Shoham S, Rottman JB, Levitz SM (2004) Involvement of CD14, toll-like receptors 2 and 4, and MyD88 in the host response to the fungal pathogen Cryptococcus neoformans in vivo. Infect Immun 72: 5373–5382.

36. Fonseca FL, Nohara LL, Cordero RJ, Frases S, Casadevall A, et al. (2010) Immunomodulatory effects of serotype B glucuronoxylomannan from Cryptococcus gattii correlate with polysaccharide diameter. Infect Immun 78: 3861–3870.

37. Nakamura K, Miyagi K, Koguchi Y, Kinjo Y, Uezu K, et al. (2006) Limited contribution of Toll-like receptor 2 and 4 to the host response to a fungal infectious pathogen, Cryptococcus neoformans. FEMS Immunol Med Microbiol 47: 148–154.

38. Biondo C, Midiri A, Messina L, Tomasello F, Garufi G, et al. (2005) MyD88 and TLR2, but not TLR4, are required for host defense against Cryptococcus neoformans. Eur J Immunol 35: 870–878.

39. Love GL, Boyd GD, Greer DL (1985) Large Cryptococcus neoformans isolated from brain abscess. J Clin Microbiol 22: 1068–1070.

40. Bacon BE, Cherniak R, Kwon-Chung KJ, Jacobson ES (1996) Structure of the O-deacetylated glucuronoxylomannan from Cryptococcus neoformans Cap70 as determined by 2D NMR spectroscopy. Carbohydr Res 283: 95–110.

41. Brummer E (1998) Human defenses against Cryptococcus neoformans: an update. Mycopathologia 143: 121–125.

42. Wang H, Li M, Lerksuthirat T, Klein B, Wuthrich M (2015) The C-Type Lectin Receptor MCL Mediates Vaccine-Induced Immunity against Infection with Blastomyces dermatitidis. Infect Immun 84: 635–642.

43. Bi L, Gojestani S, Wu W, Hsu YM, Zhu J, et al. (2010) CARD9 mediates dectin-2-induced IkappaBalpha kinase ubiquitination leading to activation of NF-kappaB in response to stimulation by the hyphal form of Candida albicans. J Biol Chem 285: 25969–25977.

44. Wozniak KL, Levitz SM (2009) Isolation and purification of antigenic components of Cryptococcus. Methods Mol Biol 470: 71–83.

45. Chen M, Wu J, Shi S, Chen Y, Wang H, et al. (2016) Structure analysis of a heteropolysaccharide from Taraxacum mongolicum Hand.-Mazz. and anticomplementary activity of its sulfated derivatives. Carbohydr Polym 152: 241–252.

46. Richards JC, Perry MB, Kniskern PJ (1984) Structural analysis of the specific polysaccharide of Streptococcus pneumoniae type 9L (American type 49). Can J Biochem Cell Biol 62: 1309–1320.

47. Urai M, Anzai H, Ogihara J, Iwabuchi N, Harayama S, et al. (2006) Structural analysis of an extracellular polysaccharide produced by Rhodococcus rhodochrous strain S-2. Carbohydr Res 341: 766–775.

48. Wang H, Shi S, Bao B, Li X, Wang S (2015) Structure characterization of an arabinogalactan from green tea and its anti-diabetic effect. Carbohydr Polym 124: 98–108.

49. Sauer KA, Scholtes P, Karwot R, Finotto S (2006) Isolation of CD4+ T cells from murine lungs: a method to analyze ongoing immune responses in the lung. Nat Protoc 1: 2870–2875.

